# Ryanodine receptor 1 is dispensable for CD4^+^ T-cell differentiation and effector function in intestinal inflammation models

**DOI:** 10.64898/2026.01.14.699480

**Authors:** Sogol Dostiar Tabrizi, Mikolaj Nawrocki, Björn-Phillip Diercks, Tanja Bedke, Marius Böttcher, Friederike Stumme, Melina Birus, Franziska Möckl, Lola Hernandez, Nicola Gagliani, Dmitri Lodygin, Alexander Flügel, Andreas Guse, Hans-Willi Mittrücker, Samuel Huber

## Abstract

T-cell receptor signaling is necessary for the activation and differentiation of CD4⁺ T cells. Calcium (Ca^2+^) signaling is essential for this process, and the complexity of Ca^2+^ channels presents a potential therapeutic target for modulating the strength of T-cell receptor signaling and further differentiation of CD4⁺ T cells. Nicotinic acid adenine dinucleotide phosphate (NAADP) is a potent Ca^2+^-mobilizing second messenger that triggers Ca^2+^ release through ryanodine receptor 1 (RYR1) in T cells. While the molecular and biophysical properties of NAADP-induced Ca^2+^ microdomains in T cells have been thoroughly investigated, and the function of the NAADP-HN1L/JPT2-RYR1 axis has been proven in T-cell activation and proliferation, its role in intestinal inflammation *in vivo* remains to be elucidated. In this study, we generated a conditional knockout mouse with *Ryr1* deleted in αβ T cells to investigate the functional relevance of RYR1 signaling in CD4⁺ T cells. *Ryr1* deletion in CD4^+^ T cells decreased TCR-induced Ca^2+^ microdomain formation, reduced peak Ca^2+^ amplitude and delayed initial velocity of global Ca^2+^ signaling *in vitro*. However, *Ryr1* expression in CD4⁺ T cells was dispensable for their pathogenicity in murine models of intestinal inflammation. Thus, *Ryr1* expression in CD4^+^ T cells plays a redundant role in intestinal inflammation.

## Introduction

Recognition of a cognate antigen via a T-cell receptor (TCR) is a hallmark of the adaptive immune system, which initiates a complex signaling network that, depending on the composition of the inflammatory milieu, directs the immune response through T-cell differentiation ^1,2^. Not only does the presence of specific cytokines in the inflammatory milieu influence the fate of CD4⁺ T cells, but the qualitative and quantitative characteristics of TCR signals also play a crucial role in directing CD4⁺ T-cell differentiation ^3,4^. A critical component of the signaling network downstream of TCR activation is calcium (Ca^2+^) signaling ^5,6^. The intricate interplay of secondary messengers, Ca^2+^ channels, and ion pumps that regulate Ca^2+^ fluxes between cellular compartments generates spatiotemporal patterns that control essential cellular events following TCR activation. The multifaceted nature of Ca^2+^ signaling in T cells therefore presents a potentially attractive therapeutic target for fine-tuning adaptive immune responses and controlling pathological chronic inflammatory conditions ^6^.

Recognition of antigen by the TCR leads to a rapid synthesis of Ca^2+^-mobilizing secondary messengers D-*myo*-inositol 1,4,5-trisphosphate (IP_3_), nicotinic acid adenine dinucleotide phosphate (NAADP), and cyclic ADP-ribose (cADPR), which induce Ca^2+^ release from intracellular stores through specific Ca^2+^ channels ^6^. Within milliseconds following TCR activation, Ca^2+^ microdomains at plasma membrane-endoplasmic reticulum junctions are formed and culminate in the vigorous release of Ca^2+^ from the endoplasmic reticulum ^7,8^. The decrease in Ca²⁺ concentration within the lumen of the endoplasmic reticulum is sensed by stromal interaction molecules (STIM1 and STIM2), which open plasma membrane Ca^2+^ release-activated Ca^2+^ channels, allowing extracellular Ca²⁺ influx into the cytosol during the process of store-operated Ca^2+^ entry (SOCE) ^9,10^. Molecular mechanisms involved in the formation of microdomains and their functional relevance have recently been a subject of thorough investigation ^8,11^. It has been shown that the key second messenger mediating the development of Ca^2+^ microdomains upon TCR activation is NAADP ^7,12^. NAADP is synthesized within milliseconds following antigen recognition by the TCR, then NAADP binds to the NAADP receptor hematological and neurological expressed 1-like protein (HN1L)/Jupiter microtubule-associated homolog 2 (JPT2) and releases Ca^2+^ from the endoplasmic reticulum via Ryanodine receptor 1 (RYR1) ^13^. In other cell types, HN1L/JPT2 also acts on acidic vesicles via two pore channels (TPC1 and TPC2) ^14^. Importantly, it has been reported that Ca^2+^ microdomains do not solely represent a stage of the Ca^2+^ signaling cascade but also have specific physiological functions, such as in T-cell migration, as well as in the trafficking of intracellular vesicles and membrane receptors ^11,15,16^.

The functional relevance of the NAADP-HN1L/JPT2-RYR1 axis has been proven to play an important role in T-cell activation and differentiation *in vitro* and *in vivo* ^14,17–20^. Small molecule antagonists of NAADP signaling impair the activation and proliferation *in vitro* in a Ca^2+^-dependent manner ^19,20^. Moreover, antagonizing NAADP skews the T-cell differentiation into a more regulatory phenotype *in vitro* and *in vivo* ^19^. Deletion of the NAADP-forming enzyme, DUOX2, or the NAADP receptor, HN1L/JPT2, resulted in decreased global Ca^2+^ signals as well as impaired activation and cytokine synthesis in murine T cells ^14^. *In vivo* studies showed that both antagonizing NAADP with the small molecule antagonist BZ194 and inhibiting ryanodine receptors with dantrolene ameliorate the activity of the experimental autoimmune encephalomyelitis (EAE) murine model of multiple sclerosis ^17,18,21^. Studying the role of RYR1 was limited by the fact that the missense mutation of the channel is particularly important for the function of striated muscle and is therefore perinatally lethal ^22^. A recent study used fetal liver chimeras to show that the deletion of *Ryr1* in the hematopoietic cells decreases the severity of symptoms in EAE and cellular analysis of these mice showed decreased percentage of CD25^+^ and CD69^+^ encephalitogenic (activated) T cells in the spinal cord of these mice compared to wild type mice (Lécuyer et al., unpublished, manuscript enclosed). Likewise, the conditional deletion of *Ryr1* in T cells confirmed this finding (Lécuyer et al., unpublished, manuscript enclosed). In contrast, mice carrying a gain-of-function *RYR1-p.R163C* mutation develop a faster and more severe EAE course compared to wild type mice ^21^.

Likewise, Trans-Ned-19, an NAADP antagonist, promotes the production of IL-10 by effector cells, increases their suppressive capacity, and induces the trans differentiation of Th17 cells into Tr1 cells, both *in vitro* and *in vivo* ^19^. However, although the function of the NAADP-HN1L/JPT2-RYR1 axis has been proven in T-cell activation and proliferation, its role in CD4^+^ T-cell differentiation and intestinal inflammation remained to be elucidated.

Inflammatory bowel disease (IBD) is a chronic immune-mediated disease with a multifactorial etiology including genetic predisposition, environmental factors, and dysregulated T-cell response ^23–25^. Differentiation of CD4^+^ Th cells, and in particular the balance between proinflammatory Th17 and Th1 cells and suppressive Foxp3^+^ Treg and IL-10 producing Tr1 cells, determines the inflammatory activity and tissue damage ^26–30^. Ca^2+^ signaling plays a critical role in the pathogenesis of IBD. Specifically, Ca^2+^ signaling in T cells was reported to control the severity of intestinal inflammation in murine models of IBD. T-cell specific deletion of components of store-operated Ca^2+^ entry leads to decreased TCR-induced Ca^2+^ signaling in T cells, reduced production of inflammatory cytokines, as well as alleviation of tissue damage in murine models of IBD ^31–34^. Moreover, inhibitors of calcineurin, a Ca^2+^-dependent phosphatase required for activation of NFAT transcription factors, are recommended for steroid-refractory cases of ulcerative colitis ^35,36^. Given the role of the NAADP-HN1L/JPT2-RYR1 axis in T-cell function in neuroinflammation, as well as the importance of Ca^2+^ signaling in inflammatory bowel disease, we aimed here to investigate the role of RYR1 in T cells during intestinal inflammation.

## Results

### Generation and validation of αβ T-cell specific RYR1 conditional knockout mouse line

Investigation of the role of RYR1 in T-cell function using knockout mice was limited by the perinatal lethality of the missense mutation of *Ryr1* ^22^. Previous studies delivered important mechanistic insights regarding the NAADP-HN1L/JPT2-RYR1 axis using chemical NAADP and RYR antagonists, or *in vitro* generated CRISPR-Cas9 knockouts ^14,19–21,46^. Nevertheless, these methods are prone to off-target effects and do not allow for a specific evaluation of the RYR1’s role in T-cell function *in vivo*.

To overcome these limitations, we generated a *Ryr1* conditional knockout mouse line. We used a conditional ready *Ryr1*^tm1a^ line, in which the *Ryr1* allele contains a gene trap cassette (lacZ and neomycin), loxP sites and FRT sites flanking exon 8. In the first step, *Ryr1^tm1a^* mice were crossed with FLP recombinase mice in order to remove the gene trap cassette and generate a functional conditional allele (*Ryr1^tm1c^*). In the next step, the resulting *Ryr1^tm1c^* mice were crossed with *CD4^cre^* mice to generate T-cell specific *Ryr1* knock-out mice (*Ryr1^tm1d^*). These mice were then further crossed to a triple reporter mouse expressing fluorescent proteins reporting for IFN-γ ^37^, IL-17A ^29^, and Foxp3 ^38^, resulting in *CD4^cre^Ryr1^+/+^ IFN-γ^katushka^IL-17^eGFP^Foxp3^RFP^*and *CD4^cre^Ryr1^fl/fl^ IFN-γ^katushka^IL-17^eGFP^Foxp3^RFP^*mice (Figure 1A, B).

**Figure 1.**
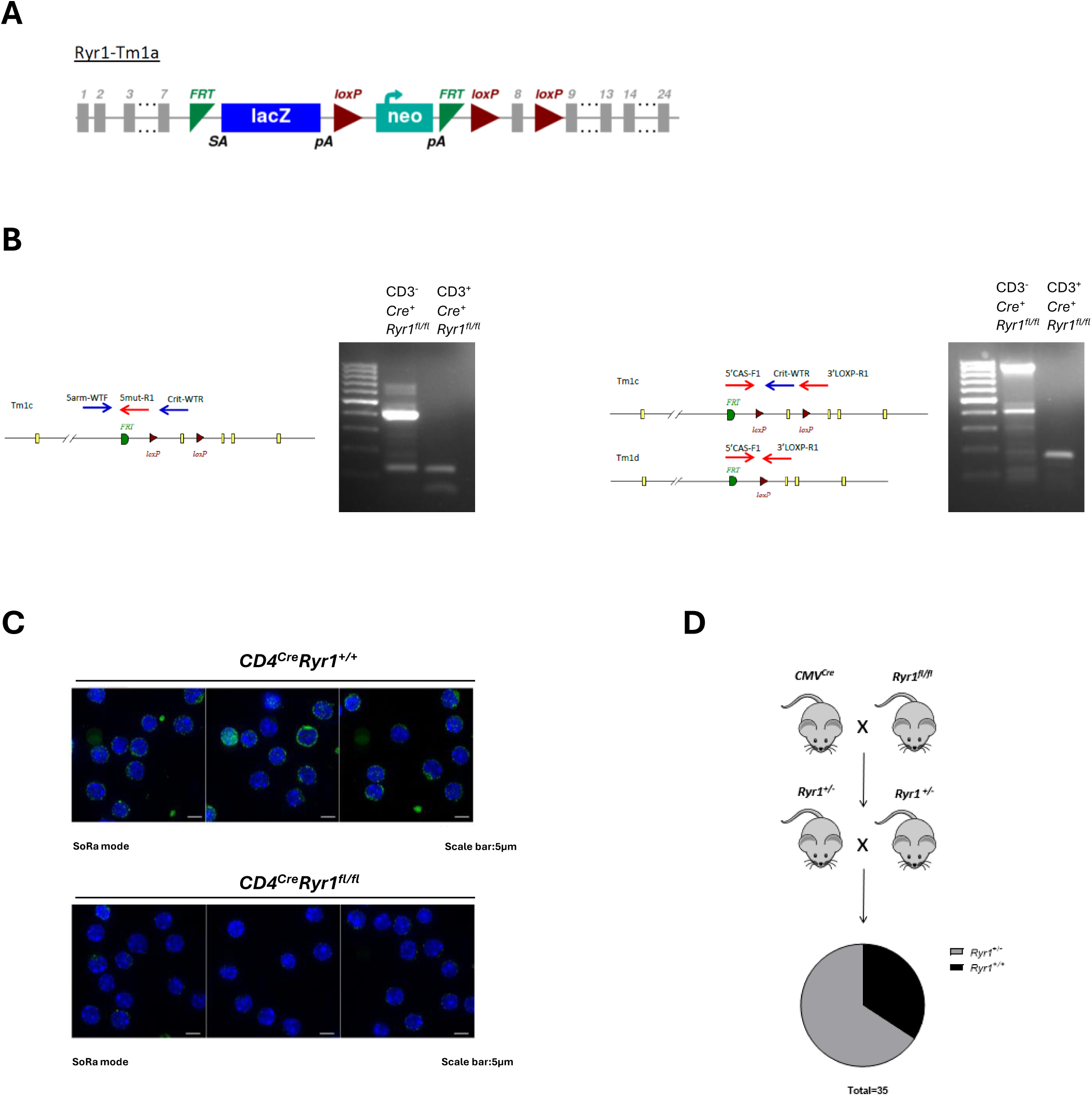
Generation and validation of the *CD4^Cre^Ryr1* conditional knockout mouse line. **A)** *Ryr1* floxed mice were generated by crossing conditional-ready *Ryr1^tm1a(EUCOMM)^* mice with a Flp recombinase-expressing strain, resulting in a *tm1c* allele with exon 8 flanked by loxP sites. **B)** Schematic of the *Ryr1^tm1c^* locus and its Cre-mediated recombination into the *Ryr1^tm1d^* allele. CD3^+^ and CD3^−^ cells were sorted by FACS from a *CD4^Cre^Ryr1^fl/fl^* mouse, and genomic DNA was analyzed by PCR to confirm the recombination. **Left**: The tm1c allele produces a 461 bp band in non-recombined CD3⁻ cells and a 125 bp mutant band in both CD3⁺ and CD3⁻ cells. **Right**: The tm1d allele yields a 174 bp product, whereas the tm1c allele generates a larger product that includes the critical region. **C)** Immunofluorescence staining of CD4^+^ T cells from *CD4^Cre^Ryr1^+/+^* and *CD4^Cre^Ryr1^fl/fl^* mice using an anti-RYR (F-1) antibody. Each field of view represents one mouse. Representative of two independent experiments. **D)** Total *Ryr1* knockout mice were generated by crossing *Ryr1^tm1c^* conditional knockout mice with a *CMV^Cre^ line*. Breeding outcomes indicate that complete loss *of Ryr1* causes embryonic or perinatal lethality.

To validate the correct integration of the conditional allele and *Ryr1* knockout efficiency, we used different sets of primers for *Ryr1^tm1c^*and *Ryr1^tm1d^*. PCR analysis of DNA from CD3^+^ and CD3^-^ cells sorted by FACS from *CD4^cre^Ryr1^fl/fl^* mice confirmed successful Cre-mediated recombination (Figure 1B). Immunofluorescence staining with a pan-RYR antibody confirmed loss of ryanodine receptors in CD4^+^ T cells in *CD4^cre^Ryr1^fl/fl^* mice on the protein level (Figure 1C). Moreover, to further validate the floxed-allele’s accuracy, we crossed *Ryr1^fl/fl^*mice with *CMV^cre^* mice and observed no viable *Ryr1^-/-^*offspring, confirming the embryonic lethality due to global *Ryr1* deletion (Figure 1D).

### Deletion of RYR1 in T cells impairs formation of Ca^2+^ microdomains and global Ca^2+^ signaling upon TCR stimulation

The endoplasmic reticulum-NAADP signaling model in T cells explains NAADP-evoked Ca^2+^ signaling. In this model, following TCR stimulation and NAADP formation, NAADP binds to its binding protein HN1L/JPT2, and the complex interacts with RYR1, leading to the formation of Ca^2+^ microdomains ^47^.

Upon TCR/CD28 stimulation, naïve CD4⁺ T cells lacking RYR1 showed a reduced number of Ca²⁺ microdomains per cell, lower Ca^2+^ signal amplitude, and decreased proportion of responder cells within the first 15 s of T-cell activation, compared to wild type controls (Figure 2A, B). To assess whether reduced Ca^2+^ microdomain formation was reflected in global cytosolic Ca²⁺ dynamics, we measured the free cytosolic Ca^2+^ concentration in naïve CD4⁺ T for 10 min after stimulation with soluble anti-CD3 mAb (Figure 2C, D). Baseline [Ca^2+^] was similar in wild type and *Ryr1-*deficient cells. However, the latency to reach the peak calcium concentration upon TCR stimulation was significantly longer in the *Ryr1*-deficient T cells compared with wild type T cells. This delay was further reflected in overall lower area under the curve in the knockout cells over a 10-minute observation period (Figure 2C, D). Thus, deletion of *Ryr1* in CD4^+^ T cells reduced TCR-induced microdomain formation, resulting in a slower rise of intracellular calcium concentration and, consequently, reduced calcium entry. These findings are consistent with previous reports using *RYR1*-knockdown Jurkat T cells ^7^, as well as results obtained with T cells derived from *Ryr1*^-/-^ fetal liver chimeric mice, which exhibited impaired microdomain formation ^7,12^.

**Figure 2.**
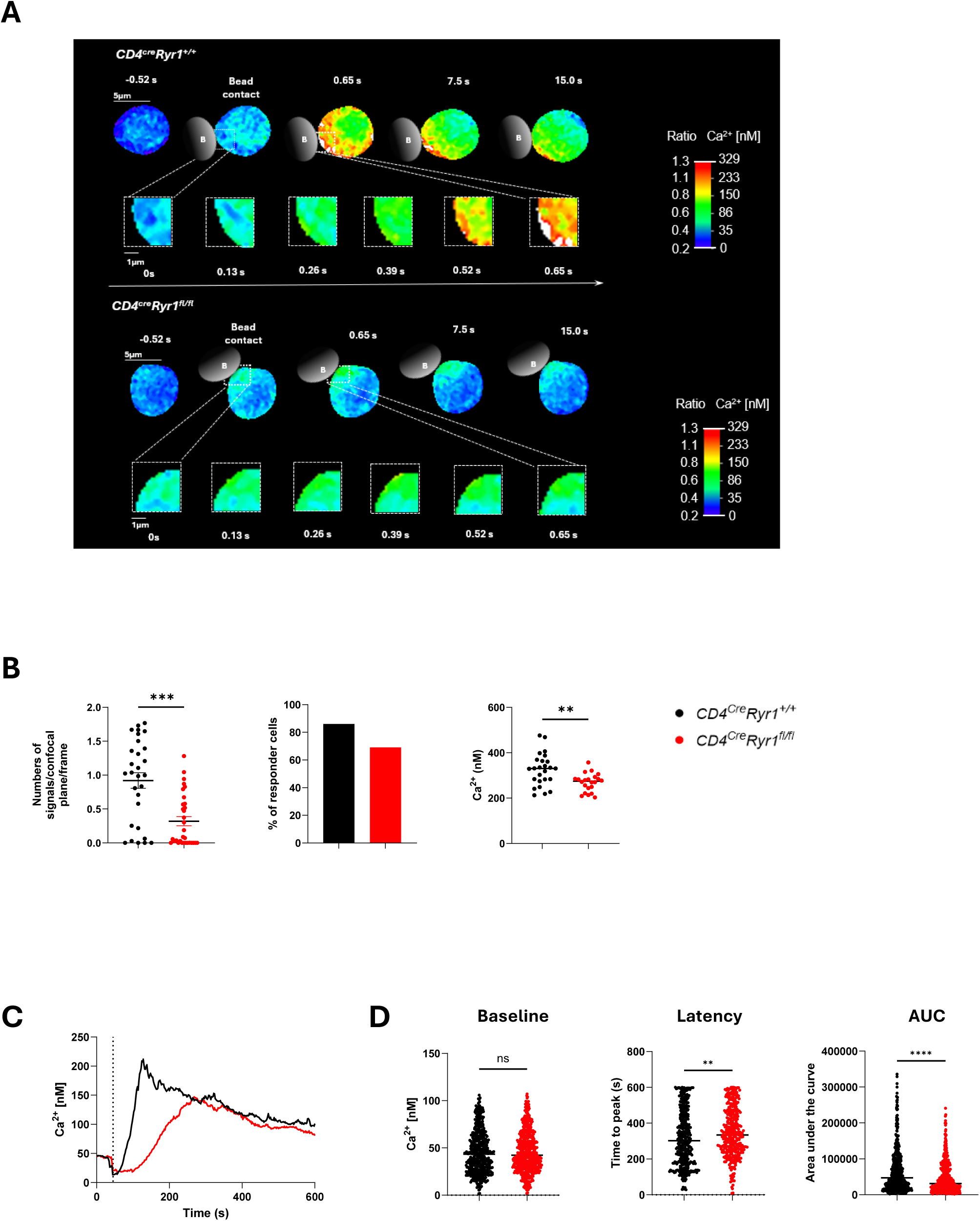
*Ryr1-*deficiency impairs early Ca²⁺ signaling in naïve CD4^+^ T cells upon TCR/CD28 stimulation. **A)** Representative images show early Ca²⁺ microdomain formation in a *CD4^cre^Ryr1^+/+^* T cell (top) and a *CD4^cre^Ryr1^fl/fl^* T cell (bottom) following stimulation with beads coated with anti-CD3 and anti-CD28 mAbs. A schematic illustrates the bead–cell contact site. Magnified views highlight regions near the bead interaction site. **B)** Quantification of Ca^2+^ microdomains formed in *CD4^Cre^Ryr1^+/+^* (n = 29) and *CD4^Cre^Ryr1^fl/fl^* (n = 32) naïve CD4^+^ T cells, in the first 15 s after contact with beads coated with anti-CD3 and anti-CD28 mAbs. Parameters analyzed include the number of signals detected per confocal plane per frame, the proportion of cells responding, and the Ca^2+^ signal amplitude. Data are presented as mean ± SEM and collected from seven independent experiments. Statistical analysis was performed using the unpaired Mann-Whitney U test. Significance: **p < 0.01, ***p < 0.001. **C)** Global Ca^2+^ signaling in *CD4^Cre^Ryr1^+/+^* (n = 496) and *CD4^Cre^Ryr1^fl/fl^* (n = 536) naïve CD4⁺ T cells. Cells were loaded with Fura-2 and stimulated with anti-CD3 mAb, followed by ratiometric Ca^2+^ imaging for 10 minutes. Data represent the mean kinetics of cytosolic Ca²⁺ concentration over time. **D)** Quantification of global Ca²⁺ responses, including baseline Ca²⁺ concentration, latency to peak, and area under the curve (AUC). Data are shown as mean ± SEM collected from 3 independent experiments. Statistical significance was determined by the unpaired Mann-Whitney U test. ns, not significant; **p < 0.01; ****p < 0.0001.

### T-cell specific deletion of *Ryr1* does not impact CD4^+^ T-cell composition in steady state

To determine whether RYR1 plays a role in T-cell development and normal immune composition, we analyzed T-cell populations and cytokine production in *CD4^cre^Ryr1^+/+^* and *CD4^cre^Ryr1^fl/fl^* mice under steady state conditions. Flow cytometry analysis of CD3^+^, CD4^+^, and CD8^+^ T cells from colon and mesenteric lymph nodes showed no significant differences in CD62L^+^CD44^low^ naïve and CD62L^-^CD44^high^ effector T-cell numbers between *CD4^cre^Ryr1^+/+^* and *CD4^cre^Ryr1^fl/fl^* mice (Figure 3A, B). In the thymus, the frequency of CD4^+^, CD8^+^, CD4^+^CD8^+^ (double positive), and CD4^-^CD8^-^ (double negative) thymocytes was comparable between groups (Figure 3C). Furthermore, Foxp3^+^ Treg cells were also present at similar frequencies in mesenteric lymph nodes and colon (Figure 3D). Upon *ex vivo* stimulation with PMA and ionomycin in the presence of monensin the production of IFN-γ, IL-17A, IL-10, and TNF-α by CD4^+^ T cells was largely comparable between wild type and *CD4^cre^Ryr1^fl/fl^*mice. TNF-α production in CD4^+^ T cells isolated from mesenteric lymph nodes was by trend higher in conditional knockout mice compared to wild type controls; nevertheless, this result did not reach statistical significance (Figure 3E). Thus, RYR1 is dispensable for the emergence of CD4^+^ T-cell subpopulations in steady state.

**Figure 3.**
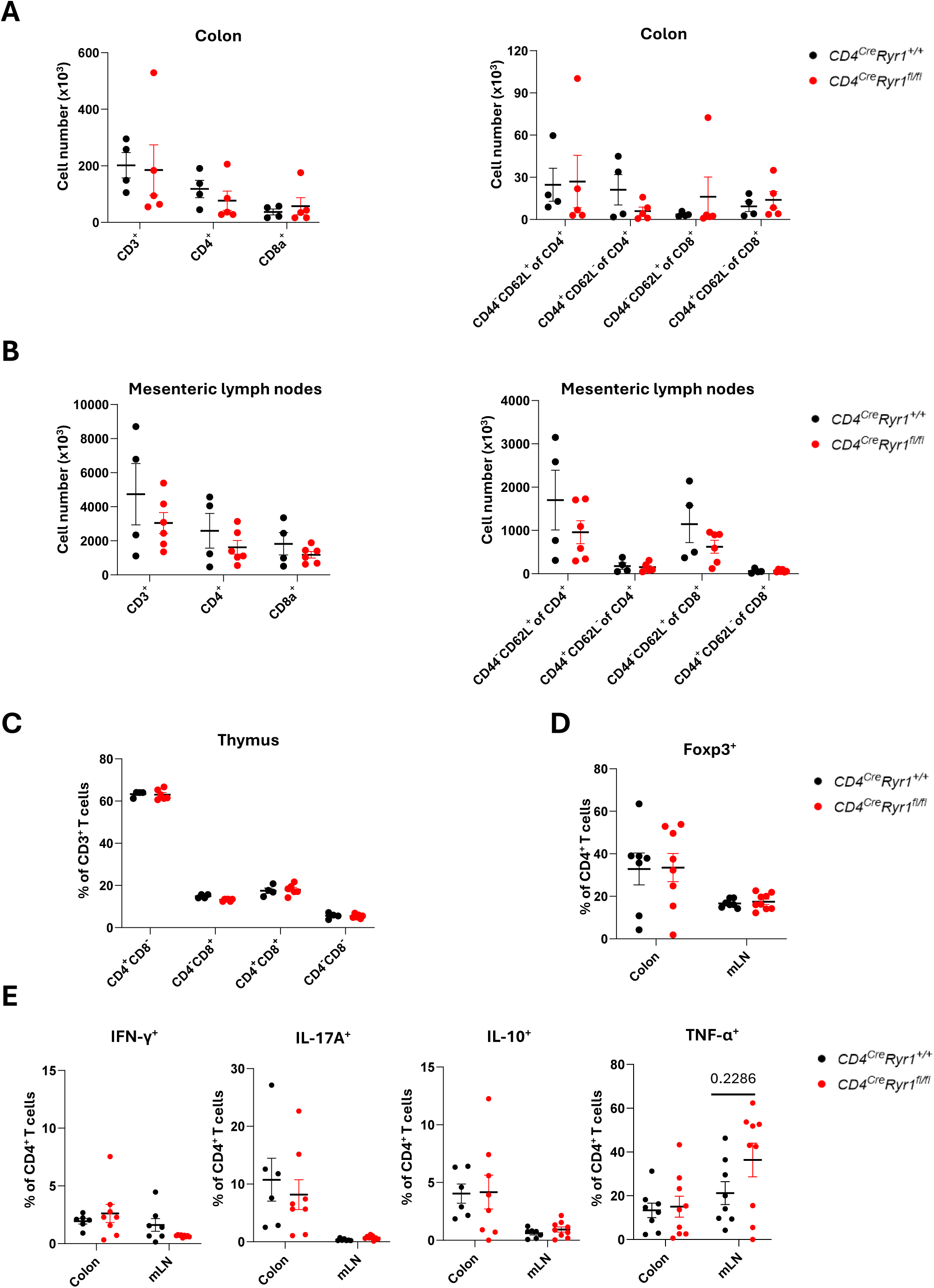
T cell compartment at steady state upon CD4Cre-driven deletion of *Ryr1*. **A, B)** Absolute numbers of CD3^+^, CD4^+^, and CD8^+^ T cells, as well as naïve (Foxp3^-^CD44^-^CD62L^+^) and effector (Foxp3^-^CD44^+^CD62L^-^) CD4^+^ and CD8^+^ T cells were measured in the **A)** colon and **B)** mesenteric lymph nodes of *CD4^Cre^Ryr1^+/+^* and *CD4^Cre^Ryr1^fl/fl^* mice (n=4-6 mice in each group, 2 independent experiments). **C)** Frequencies of CD4^+^, CD8^+^, double-positive (DP), and double-negative (DN) thymocytes in *CD4^Cre^Ryr1^+/+^* and *CD4^Cre^Ryr1^fl/fl^* mice (n=4-6 mice in each group, 2 independent experiments). **D)** Frequencies of Tregs in the colon and mesenteric lymph nodes of *CD4^Cre^Ryr1^+/+^* and *CD4^Cre^Ryr1^fl/fl^* mice (n=7-8 mice in each group, 3 independent experiments). **E)** Cytokine production (IFN-γ, IL-17A, TNF-α, and IL-10) by CD4^+^ T cells isolated from the colon and the mesenteric lymph nodes of *CD4^Cre^Ryr1^+/+^* and *CD4^Cre^Ryr1^fl/fl^* mice following stimulation with PMA and ionomycin (n=6-9 mice in each group, 3 independent experiments). Mice were 7-23 weeks old at the time of analysis. All data are presented as the mean ± SEM. Statistical comparisons were performed using the unpaired Mann-Whitney U test.

### *Ryr1* expression is higher in effector subsets of CD4^+^ T cells

Next, we sought to determine the expression pattern of *Ryr1* in CD4⁺ T-cell subsets. To this end, we sorted Foxp3⁺ Treg cells, CD62L⁺CD44^low^ naïve, and CD62L⁻CD44^high^ effector CD4⁺ T cells from the spleens of wild type mice under steady-state conditions by FACS. Quantitative RT-PCR analysis revealed that *Ryr1* expression was higher in effector T cells compared to naïve and regulatory T cells (Figure 4A). To further investigate *Ryr1* expression in effector subsets and Foxp3⁺ Treg cells, during intestinal inflammation, we induced transient intestinal inflammation by injecting anti-CD3 mAb into *IFN-γ^katushka^ IL-17^eGFP^ Foxp3^RFP^* reporter mice, which express fluorescent proteins upon Foxp3, IL-17A, or IFN-γ expression. Using flow cytometry, we sorted IFN-γ^Katushka+^ Th1, IL-17A^eGFP+^ Th17, and IFN-γ^Katushka+^ IL-17A^eGFP+^ Th1/Th17 cells, as well as Foxp3⁺ Treg cells, and quantified *Ryr1* expression in these subsets. Among these subsets, *Ryr1* expression was specifically elevated in Th1 cells (Figure 4B). Taken together, *Ryr1* is preferentially expressed in effector CD4^+^ Th1 cells.

**Figure 4.**
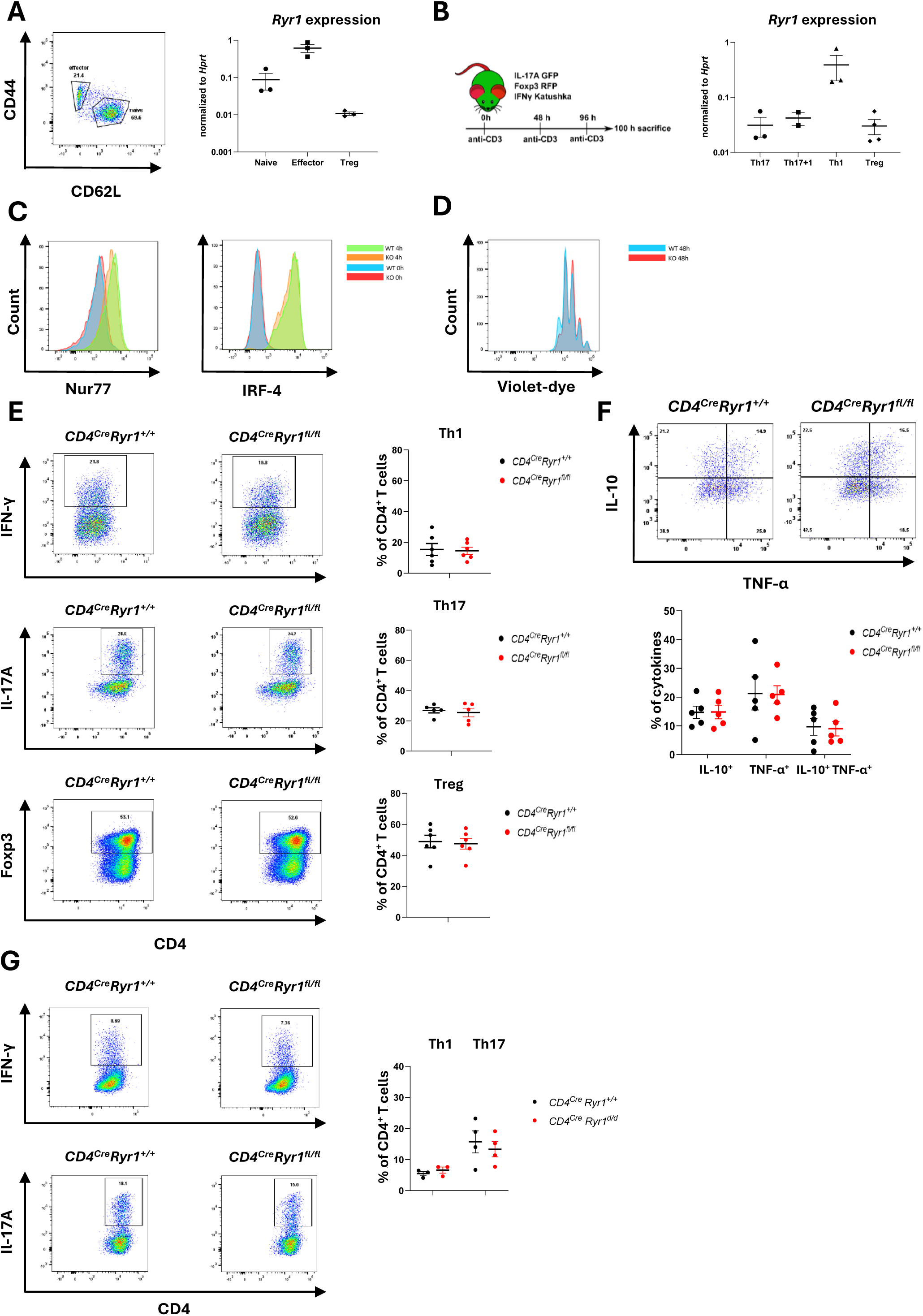
*Ryr1* expression and functional characterization in CD4^+^ T Cell activation, proliferation, and *in vitro* differentiation. **A, B)** Quantitative RT-PCR analysis of *Ryr1* expression in **A)** CD62L^+^CD44^low^ naïve and CD62L^-^CD44^high^ effector CD4^+^ T cells, and Foxp3^+^ T regulatory cells (Tregs) from spleens of WT mice at steady state (3 biological replicates per group), and **B)** in Th1, Th17, Th1+17, and Tregs sorted using FACS from small intestines of *Foxp3^RFP^ IL-17A^eGFP^ IFN-γ^Katushka^* triple reporter mice treated after induction of intestinal inflammation by anti-CD3 mAb administration (n = 2–4 biological replicates per group). **C)** Naïve CD4^+^ T cells from *CD4^cre^Ryr1^+/+^* and *CD4^cre^Ryr1^fl/fl^* mice were restimulated *in vitro* with anti-CD3 and anti-CD28 mAbs. Expression of early TCR activation markers Nur77 and IRF4 was measured by FACS at 4 and 16 hours post-restimulation, respectively (n = 3 biological replicates per group). **D)** Naïve CD4^+^ T cells from *CD4^cre^Ryr1^+/+^* and *CD4^cre^Ryr1^fl/fl^* mice were labeled with CellTrace Violet before restimulation with anti-CD3 and anti-CD28 mAbs. Proliferation was assessed by CellTrace Violet dye dilution via flow cytometry after 48 hours (n = 3 biological replicates per group). **E)** *In vitro* differentiation of naïve CD4^+^ T cells into Th1, Th17, and Treg. FACS sorted CD62L⁺CD44^low^ CD4⁺ T cells from *CD4^cre^Ryr1^+/+^* and *CD4^cre^Ryr1^fl/fl^* mice were cultured for 96 hours under polarizing conditions. Cytokine expression was assessed after 4 hours of restimulation with ionomycin and PMA. Shown are representative dot plots and frequencies of IFN-γ^+^ Th1 cells, IL-17A^+^ Th17 cells, and Foxp3^+^ Treg cells (n = 6 biological replicates per group). **F)** *In vitro* restimulation of effector CD4^+^ T cells. FACS sorted CD62L⁻CD44^high^ effector CD4⁺ T cells from *CD4^cre^Ryr1^+/+^* and *CD4^cre^Ryr1^fl/fl^* mice were cultured with anti-CD3 and anti-CD28 mAbs in the presence of IL-2 for 96 hours. Cytokine expression was assessed after 4 hours of restimulation with ionomycin and PMA. Shown are representative dot plots and frequencies of effector CD4^+^ T cells producing IL-10 and TNF-α (n = 5 biological replicates per group). **G)** *In vitro* differentiation of naïve CD4^+^ T cells into Th1, Th17, and Treg. FACS sorted CD62L⁺CD44^low^ CD4⁺ T cells from *CD4^cre^Ryr1^+/+^IL-17A^eGFP^IFN-γ^Katushka^* and *CD4^cre^Ryr1^fl/fl^ IL-17A^eGFP^IFN-γ^Katushka^* mice were cultured for 96 hours under polarizing conditions. Cytokine expression was assessed without restimulation. Shown are representative dot plots and frequencies of IFN-γ^+^ Th1 cells, IL-17A^+^ Th17 cells, and Foxp3^+^ Treg cells (n = 3–4 biological replicates per group). All experiments were independently performed at least three times. All data are presented as the mean ± SEM. Statistical comparisons were made using the unpaired Mann-Whitney U test.

### *Ryr1*-deficiency does not impair CD4^+^ T-cell activation and differentiation *in vitro*

Ca^2+^ signaling is essential for regulating primary T-cell activation, function, cytokine production, and proliferation ^6,48–51^ and disruption of Ca^2+^ signaling impairs T-cell response and function ^32^. Therefore, we sought to investigate whether RYR1-mediated Ca²⁺ flux plays a role in the activation and differentiation of CD4⁺ T cells. Therefore, we next analyzed early activation markers in naïve CD4^+^ T cells following TCR stimulation. To this end, we isolated naïve CD4^+^ T cells from *CD4^cre^Ryr1^+/+^* and *CD4^cre^Ryr1^fl/fl^* mice and stimulated them *in vitro* with anti-CD3 and anti-CD28 mAbs. Transcription factors Nur77 and IRF4 are markers of T-cell activation whose expression is correlated with TCR signal strength ^52,53^. No significant difference in the upregulation of these markers was observed between *CD4^cre^Ryr1^+/+^*and *CD4^cre^Ryr1^fl/fl^* T cells (Figure 4C, D). In line with these findings, *Ryr1*-deficiency did not change T-cell proliferation under *in vitro* conditions (Figure 4E). These results suggest that RYR1 is dispensable for early TCR signaling and initial T-cell activation. To determine whether RYR1 affects CD4⁺ T-cell differentiation and cytokine production, we sorted CD62L⁺CD44^low^ naïve and CD62L⁻CD44^high^ effector CD4⁺ T cells by FACS from *CD4^cre^Ryr1*^+/+^ and *CD4^cre^Ryr1*^fl/fl^ mice and stimulated them *in vitro*. Naïve cells were polyclonally activated using anti-CD3 and anti-CD28 mAbs and cultured under Th1 (IL-12, IL-2, and anti–IL-4 mAb), Th17 (IL-6, TGF-β, anti–IL-4 mAb, and anti–IFN-γ mAb), and Treg (TGF-β, IL-2) polarizing conditions to promote subset-specific differentiation. Effector T cells were restimulated with anti-CD3 and anti-CD28 mAbs in the presence of IL-2, without additional polarizing cytokines, to assess their cytokine production capacity. Cytokine production and transcription factor expression were assessed after 4 days using flow cytometry. For intracellular cytokine staining, cells were restimulated with PMA, ionomycin, and monensin. Flow cytometric analysis revealed no differences in *in vitro* differentiation of naïve T cells or cytokine production by effector T cells (Figure 4F, G).

Since PMA and ionomycin bypass physiological Ca^2+^ signaling and may override the effects of impaired RYR1-mediated Ca^2+^ release, we repeated the assays without restimulation. We crossed the *CD4^cre^Ryr1^fl/fl^*line with reporter mice expressing fluorescent proteins for IL-17A and IFN-γ. In line with previous findings, *Ryr1*-deficiency did not alter IL-17A or IFN-γ expression under Th17 and Th1 polarizing conditions, respectively (Figure 4H). These results indicate that RYR1 is not essential for CD4⁺ T-cell differentiation or cytokine production *in vitro*.

In summary, these results show that *Ryr1* expression is upregulated in effector T cells, particularly in Th1 cells. However, *Ryr1*-deficiency did not change early TCR activation, T-cell proliferation, differentiation, and cytokine production *in vitro*.

### *Ryr1*-deficiency does not significantly alter T-cell mediated acute and chronic intestinal inflammation

Our *in vitro* data indicated that RYR1 is dispensable for the signaling cascade downstream of TCR activation as well as for T-cell proliferation and differentiation into Th1, Th17, and Foxp3^+^ Treg cells. Nevertheless, *Ryr1* expression is upregulated in effector T cells, while naïve T cells express low amounts. Furthermore, *in vitro* assays cannot fully reproduce the complexity of inflammatory responses *in vivo*. Therefore, we next aimed to investigate whether RYR1 signaling in T cells plays a role *in vivo* in the context of intestinal inflammation. Indeed, mucosal immunity is regulated by a balance between effector Th1 and Th17 cells, as well as regulatory Foxp3^+^ T cells and Foxp3^-^ T regulatory 1 (Tr1) cells, and is shaped by complex interactions between immune cells, microbiota-derived signals, cytokine networks, and environmental factors ^54^.

To evaluate the role of RYR1 signaling in T cells during acute intestinal inflammation, *CD4^cre^Ryr1^+/+^* and *CD4^cre^Ryr1^fl/fl^*mice were administered anti-CD3 mAb systemically. This model is characterized by systemic T-cell activation followed by global inflammatory responses with a cytokine storm. These responses promote Th17-cell differentiation and their migration to small intestine as well as expansion of Foxp3^+^ Treg cells and Tr1 cells which suppress Th17 cells via IL-10 signaling ^29,30^. Repeated anti-CD3 mAb injections result in intestinal inflammation, that primarily affects the upper small intestine and causes significant weight loss recapitulating acute immune-mediated tissue damage ^30,55^.

Following anti-CD3 mAb injection, both *CD4^cre^Ryr1^+/+^* and *CD4^cre^Ryr1^fl/fl^* mice exhibited rapid weight loss due to systemic inflammatory response and diarrhea. The extent of the weight loss was comparable between the groups, with an average of 15%, revealing similar levels of inflammation in both genotypes (Figure 5A, B). In line with these data, flow cytometric analysis of mesenteric lymph nodes showed no significant differences in the frequencies and absolute cell numbers of Th1, Th17, and Foxp3^+^ Treg cells (Figures 5C, D).

**Figure 5.**
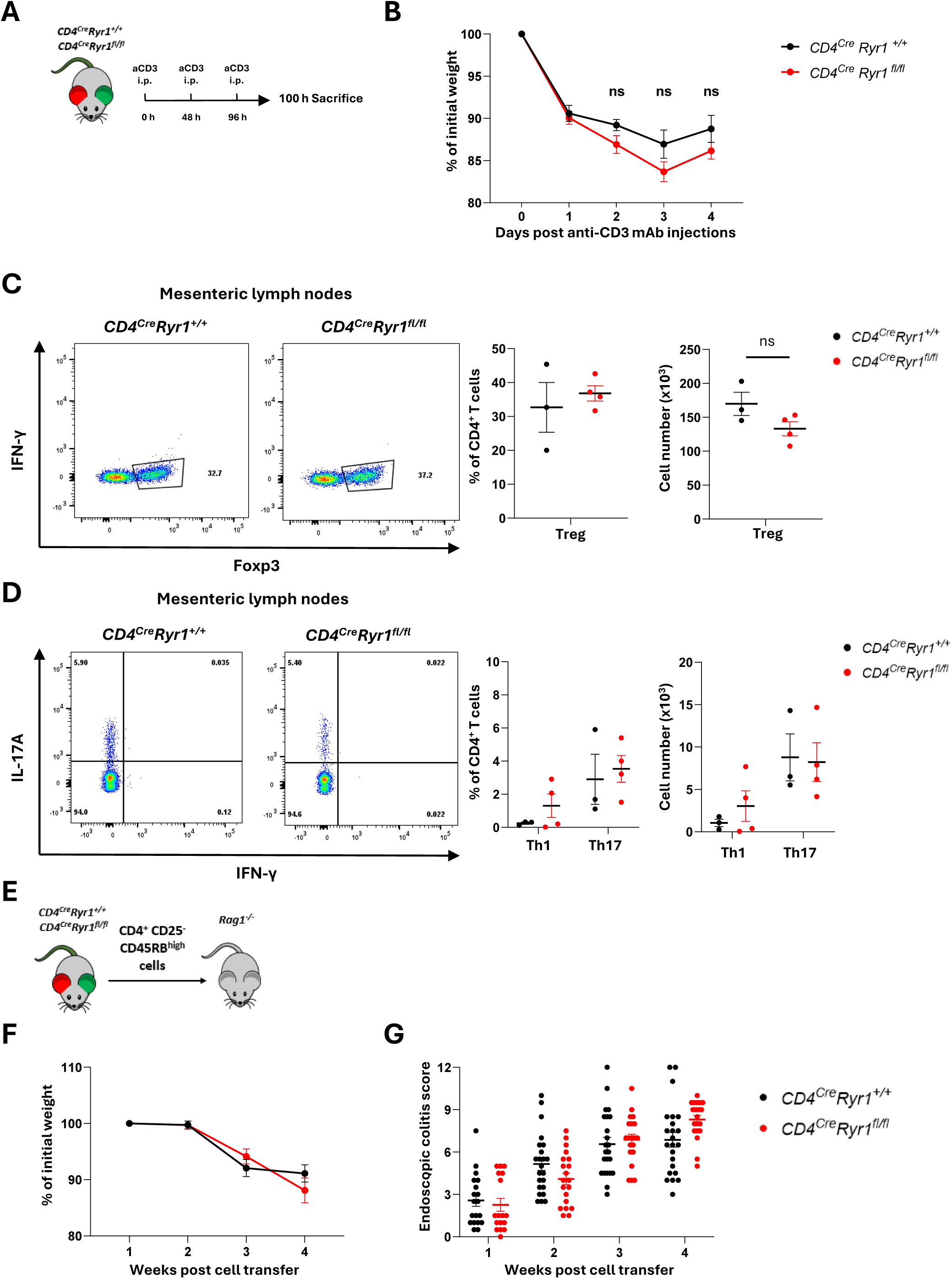
T-cell specific deletion of *Ryr1* does not alter T-cell mediated acute or chronic intestinal inflammation. **A)** Schematic of anti-CD3 mAb-induced intestinal inflammation model. *CD4^Cre^Ryr1^+/+^* and *CD4^Cre^Ryr1^fl/fl^* mice were injected with anti-CD3 mAb on days 0, 2, and 4, and the mice were sacrificed 4 hours after the final injection. **B)** Weight loss curves in *CD4^Cre^Ryr1^+/+^* and *CD4^Cre^Ryr1^fl/fl^* mice during anti-CD3 mAb-induced small intestinal inflammation (n= 6-7 mice in each group, two independent experiments). **C, D)** Representative dot plots and quantification of **C)** Foxp3^+^ Treg cell frequency and absolute number, and **D)** IFN-γ^+^ Th1 and IL-17A^+^ Th17 cell frequencies and absolute numbers in mesenteric lymph nodes from anti-CD3 mAb-treated *CD4^Cre^Ryr1^+/+^* and *CD4^Cre^Ryr1^fl/fl^* mice after 4 hours of restimulation with PMA and ionomycin (n=3-4 mice in each group, one experiment). **E)** Schematic of the adoptive transfer colitis model. FACS-sorted CD45RB^high^CD4^+^ T cells from *CD4^cre^Ryr1^+/+^* and *CD4^cre^Ryr1^fl/fl^* mice were transferred to *Rag1^-/-^* recipient mice (n=43 total, 20-23 mice in each group, 3 independent experiments). **F, G)** Weight loss and colonoscopy colitis scores of *Rag1^-/-^* mice following CD45RB^high^CD4^+^ T cell transfer. All data are presented as the mean ± SEM. Two-way repeated measures ANOVA with Bonferroni’s post hoc test was used for weight loss and colonoscopy scores **(B, F, G),** and the Mann-Whitney U test was used for flow cytometry comparisons **(C, D)**. ns = not significant. Each dot represents an individual mouse.

Since anti-CD3 mAb administration induces acute inflammation, this model is more suitable for studying T-cell activation and differentiation *in vivo* but does not mimic the chronic inflammation observed in IBD. Moreover, *Ryr1* expression is elevated in effector T cells, and the short duration of the anti-CD3 mAb model may not sufficiently capture the impact of *Ryr1* deletion on their function.

To address this, we employed an adoptive transfer colitis model, which is characterized by chronic intestinal inflammation. In this system, naïve CD45RB^high^CD4^+^ T cells are transferred into *Rag1^-/-^*mice, which lack functional T and B cells. Upon transfer, the cells predominantly differentiate into Th1 and Th17 effector cells, while conversion into Treg cells is limited. This imbalance makes the model a valuable tool to study the immunopathogenesis of IBD in mice ^56,57^. Regardless of the *Ryr1* genotype of donor T cells, both groups developed colitis with similar colitis scores and weight loss, showing that *Ryr1* expression in CD4^+^ T cells is dispensable for disease progression and severity in this chronic inflammatory model (Figure 5E-G).

### *Ryr1* expression in Th1 and Th17 cells is dispensable for T-cell-mediated colitis progression

While the transfer of CD45RB^high^CD4^+^ T cells mimics the chronic T-cell driven colitis through the activation and differentiation of naïve T cells, it does not allow for a direct assessment of the RYR1’s role in specific effector T-cell subsets, where its expression is more pronounced. Given that *Ryr1* expression is higher in effector T cells compared to naïve T cells, especially in Th1 cells, and that both Th1 and Th17 cells are important mediators of inflammatory bowel diseases ^58–60^, we next focused on these T-cell subsets.

To assess whether *Ryr1* deficiency in Th1 and Th17 cells influences colitis pathogenesis, we used a model of effector T-cell transfer colitis. *CD4^cre^Ryr1^+/+^* and *CD4^cre^Ryr1^fl/fl^*mice backcrossed to the *IFN-γ^katushka^IL-17^eGFP^Foxp3^RFP^*reporter line were used for these experiments.

To examine the role of RYR1 in Th1 cells, we co-transferred FACS-sorted *in vitro* differentiated wild type Foxp3^-^ Th17 cells in combination with either wild type or *Ryr1* deficient Foxp3^-^ Th1 cells into *Rag1^-/-^* recipient mice (Figure 6A). This approach ensured robust colitis induction while specifically testing the role of RYR1-mediated Ca^2+^ signaling in transferred Th1 cells. Colitis developed 1-2 weeks after cell transfer. Colitis severity, assessed by weight loss and weekly endoscopic scoring, revealed that the presence or absence of RYR1 in Th1 cells did not alter the pathogenicity of Th17 cells (Figure 6B-D). These findings indicate that *Ryr1* deficiency in transferred Th1 cells does not change the progression or severity of colitis when co-transferred with wild type Th17 cells, suggesting that RYR1 is dispensable for the function of transferred Th1 cells in this model.

**Figure 6.**
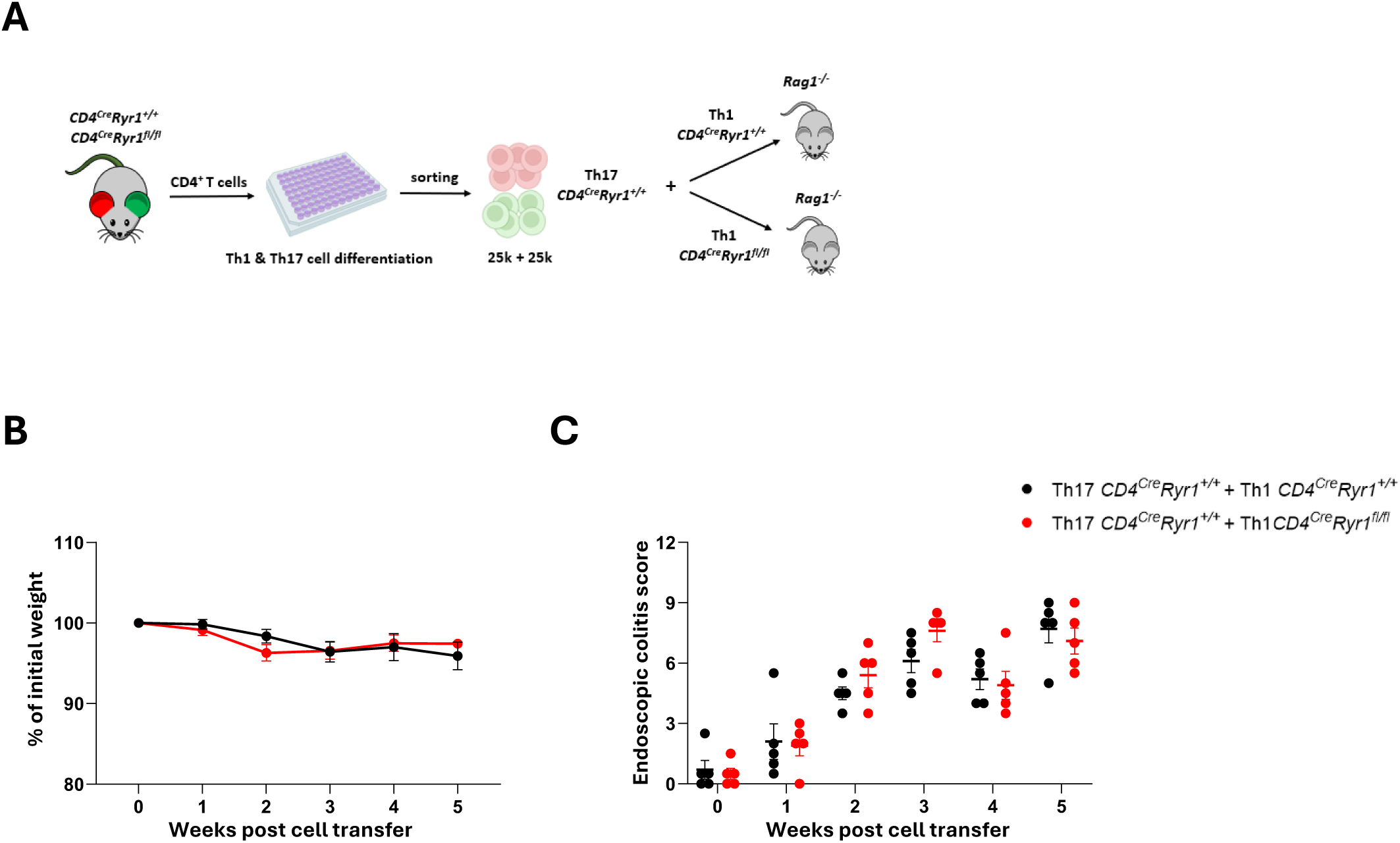
*Ryr1* absence in Th1 cells does not affect colon inflammation severity in effector cell transfer colitis. **A)** Schematic representation of Th17 + Th1 cell co-transfer colitis. CD4^+^ T cells were isolated from *CD4^Cre^Ryr1^+/+^* and *CD4^Cre^Ryr1^fl/fl^ Foxp3^RFP^IL-17A^eGFP^IFN-γ^Katushka^* triple reporter mice. CD4^+^ T cells from *CD4^Cre^Ryr1^+/+^* mice were differentiated *in vitro* into Th1 and Th17 cells. CD4^+^ T cells from *CD4^Cre^Ryr1^fl/fl^* mice were differentiated exclusively into Th1 cells. Th1 (IFN-γ^+^) and Th17 (IL-17A^+^) cells were sorted by FACS. A total of 2.5×10^4^ *CD4^Cre^Ryr1^+/+^* Th17 combined with 2.5×10^4^ Th1 cells, either from *CD4^Cre^Ryr1^+/+^* or *CD4^Cre^Ryr1^fl/fl^* mice, were transferred into *Rag1^-/-^* recipient mice (n=10 total, 5 in each group, one experiment). **B)** Weekly weight loss and **C)** endoscopic colitis scores. All data are presented as the mean ± SEM. Statistical comparisons were made using the two-way repeated measures ANOVA with Bonferroni’s post hoc test. Each dot represents an individual mouse.

To assess how RYR1 influences the pathogenicity of Th17 cells as well as their trans differentiation into Th1 cells, we transferred *in vitro* differentiated Foxp3^-^ Th17 cells either from *CD4^cre^Ryr1^+/+^*or from *CD4^cre^Ryr1^fl/fl^* mice into *Rag1^-/-^* recipient mice (Figure 7A). Both wild type Th17 cells and *Ryr1*-deficient Th17 cells induced similar levels of intestinal inflammation, as measured by weight loss and endoscopic score (Figures 7B, C). Flow cytometry analysis of T cells isolated from colon and mesenteric lymph nodes showed that a small proportion of T cells were still expressing IL-17A, and a similar proportion expressing both IL-17A and IFN-γ, while the majority of T cells expressed IFN-γ only, suggesting that a major proportion of transferred Th17 cells trans differentiated into Th1 cells (Figure 7D, E). In the draining lymph nodes, the frequency of Th1 cells was lower in the mice that received *Ryr1*-deficient cells compared to wild type controls; however, this difference did not reach statistical significance. No significant differences were observed among T-cell subset frequencies, and absolute cell numbers between wild type and *Ryr1* deficient Th17 cells. These data indicate that RYR1-mediated Ca^2+^ signaling is dispensable for Th17 cell-driven colitis in this model.

**Figure 7.**
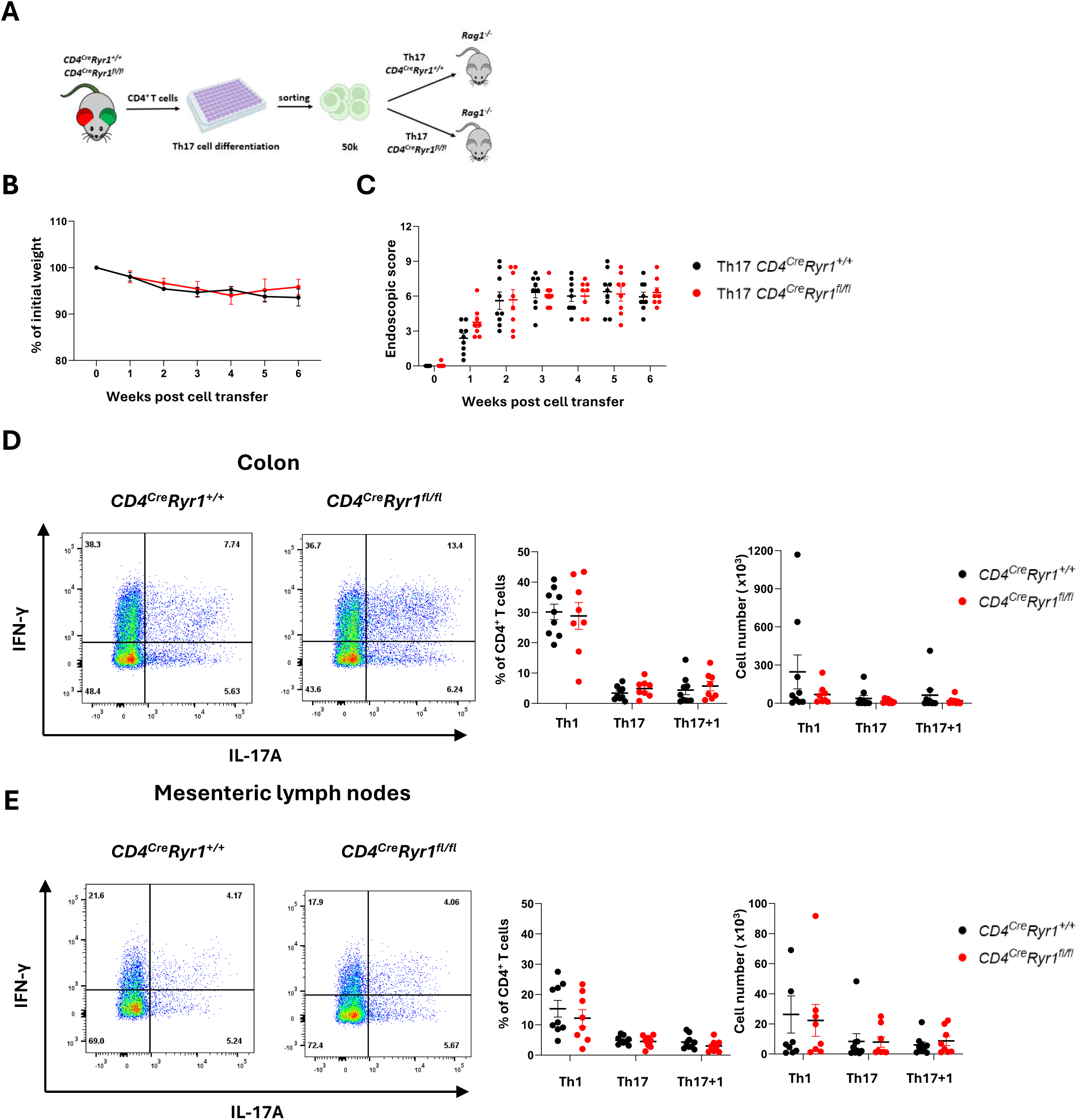
Ryr1 deficiency in Th17 cells does not affect disease severity in a Th17 adoptive transfer model of colitis. **A)** Schematic representation of the Th17 cell transfer colitis model. CD4^+^ T cells were isolated from *CD4^Cre^Ryr1^+/+^* and *CD4^Cre^Ryr1^fl/fl^ Foxp3^RFP^IL-17A^eGFP^IFN-γ^Katushka^* triple reporter mice and differentiated *in vitro* into Th17 cells. Th17 (IL-17A^+^) cells were sorted by FACS and 5×10^4^ of either *CD4^Cre^Ryr1^+/+^* or *CD4^Cre^Ryr1^fl/fl^* Th17 cells were transferred into *Rag1*^-/-^ mice (n=17 total, 8-9 mice per group, two independent experiments). **B)** Weight loss and **C)** endoscopic colitis scores at indicated time points. **D, E)** Representative dot plots and quantification of Th1, Th17, and Th1+Th17 double-positive cells in the **D)** colon and **E)** mesenteric lymph nodes shown as both frequency and absolute number. Mice were sacrificed at week 6 after transfer. All data are presented as the mean ± SEM. Two-way repeated measures ANOVA with Bonferroni’s post hoc test was used for weight loss and colonoscopy scores **(B, C)**; Mann–Whitney U test was used for flow cytometry comparisons **(D, E)**. Each dot represents an individual mouse.

To further investigate the role of RYR1 in colitis mediated by both Th1 and Th17, *in vitro* polarized wild type and *Ryr1*-deficient Th1 and Th17 cells were sorted by flow cytometry and transferred into *Rag1^-/-^*recipient mice (Figure 8A). Mice developed signs of colitis such as colon inflammation, diarrhea, and weight loss within 2–3 weeks (Figure 8B, C). While there were no significant differences in weight loss and colitis score, we observed a trend towards reduced disease severity in *Rag1^-/-^* reconstituted with *Ryr1*-deficient Th1 and Th17 cells compared to control, both in the weight loss as well as the endoscopic colitis score (Figure 8B, C). Flow cytometry analysis of the inflamed colons showed trends towards higher frequency of *CD4^cre^Ryr1^fl/fl^*Th1 cells, as well as trends for decreased numbers of *CD4^cre^Ryr1^fl/fl^*Th1 and Th17 cells. However, these findings did not reach statistical significance. The percentages and numbers of effector cell subsets were comparable in the mesenteric lymph nodes (Figure 8D, E).

**Figure 8.**
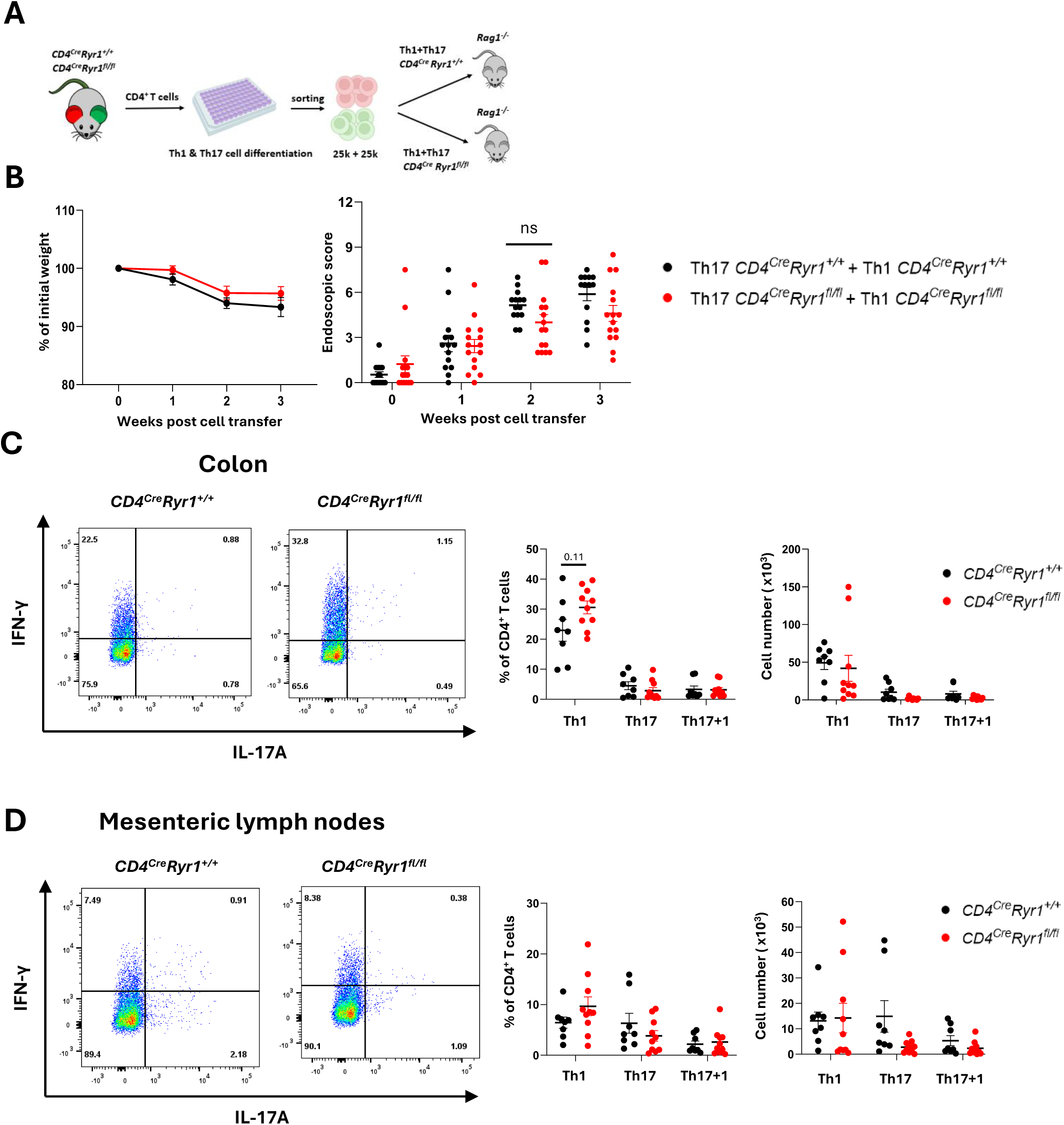
Effect of *Ryr1*-deficiency in Th1 and Th17 cells on disease severity in an adoptive transfer colitis model. **A)** Schematic representation of the experimental setup. CD4^+^ T cells were isolated from *CD4^Cre^Ryr1^+/+^* and *CD4^Cre^Ryr1^fl/fl^ Foxp3^RFP^IL-17A^eGFP^IFN-γ^Katushka^* triple reporter mice and differentiated *in vitro* into Th1 and Th17 cells. 2.5×10^4^ of FACS sorted Th1 (IFN-γ+) and Th17 (IL-17A+) cells *were* co-transferred into *Rag1*^-/-^mice (n=29 total, 14-15 mice per group, four independent experiments). **B)** Weight loss and endoscopic colitis scores over the course of the disease (data from four independent experiments). **C and D)** Flow cytometric analysis of cells isolated from the colonic mucosa and mesenteric colon-draining lymph nodes. Shown are representative dot plots and quantification of absolute cell counts and frequencies of Th1, Th17, Th1+Th17 double-positive, and Tregs (data from three independent experiments). All data are presented as the mean ± SEM. Two-way repeated measures ANOVA with Bonferroni’s post hoc test was used for weight loss and colonoscopy scores **(B)**; the Mann-Whitney U test was used for flow cytometry comparisons **(C, D)**. Each dot represents an individual mouse.

Taken together, these results indicate that *Ryr1* deficiency in Th1 and Th17 does not significantly impact colitis development and severity. Thus, RYR1-mediated Ca^2+^ signaling in Th17 and Th1 cells seems to play a redundant function in intestinal inflammation.

## Discussion

Ryanodine receptor 1 has been identified as a key mediator in the formation of Ca²⁺ microdomains within milliseconds of TCR stimulation in CD4⁺ T cells. Previous studies have demonstrated that pharmacological inhibition of ryanodine receptors or disruption of NAADP-mediated Ca^2+^ signaling impairs CD4⁺ T-cell activation, proliferation, and differentiation ^14,17–20^. In this study, we generated and validated a *CD4^cre^Ryr1^fl/fl^* conditional knockout mouse line, in which *Ryr1* is deleted in αβ T cells. Single-cell imaging of intracellular Ca^2+^ dynamics following TCR stimulation confirmed the critical role of RYR1 in the formation of Ca^2+^ microdomains within seconds after TCR stimulation. A lower number of initial Ca^2+^ microdomains in *Ryr1-*deficient cells led to a longer latency to peak Ca^2+^ levels and consequently to reduced overall Ca^2+^ influx. These results confirmed the findings obtained with Jurkat cells with knockdown of *RYR1* and with *Ryr1^-/-^* murine T cells from fetal liver chimeras ^7,12^. Interestingly, altered Ca^2+^ signaling did not affect the proliferation or differentiation of CD4^+^ T cells *in vitro* and *in vivo upon* polyclonal stimulation. It remains unclear whether TCR signal strength functions as a continuous variable that is proportionally translated into cell fate, or whether it is discretized into several thresholds at the single-cell level. Increasing evidence supports the latter^61,62^. Our findings suggest that Ca^2+^ may follow the same threshold-dependent mechanism, and the single knockout of *Ryr1* does not impair TCR-dependent calcium signaling to a degree that affects cell proliferation or differentiation.

Early models proposed that TCR-dependent Ca^2+^ signaling is initiated by the formation of localized Ca^2+^ signals generated through NAADP-mediated release of Ca^2+^ from the endoplasmic reticulum via RYR1^7^. Recent evidence has revealed the complexity of the NAADP signaling network, identifying two-pore channels 1 and 2, along with accessory binding proteins, as its additional components. Specifically, NAADP binding to HN1L/JPT2 facilitated Ca²⁺ release via both RYR1 and TPC1, while its interaction with LSM12 primarily promotes Ca²⁺ release through TPC2. Nevertheless, these broader models suggest that NAADP functions as a versatile second messenger that regulates Ca²⁺ signaling through multiple channels, providing redundancy among specific calcium channels in T cells and enabling possible compensatory mechanisms within this pathway ^47,63,64^. Moreover, ryanodine receptors, like IP_3_ receptors, are activated by Ca^2+^ through the process of calcium-induced calcium release ^65^. Studies in skeletal muscle cells isolated from myopathy patients with impaired RYR1 function and mouse muscle cells in which RYR1 levels were reduced using siRNA have shown that impaired RYR1 function increased IP_3_Rs expression ^66^. Whether similar compensatory mechanisms occur in T cells remains to be investigated. Additionally, it is important to consider that antagonists of ryanodine receptors or of NAADP-mediated Ca^2+^ signaling act within hours, leading to impaired T-cell activation and proliferation and potentially not allowing sufficient time for cells to activate compensatory pathways. In contrast, in the case of the conditional deletion of *Ryr1*, cells might upregulate alternative Ca^2+^ signaling pathways or channels, thereby compensating for the absence of RYR1 and preserving normal activation and proliferative capacity.

Ryanodine receptor 1 plays a critical role in striated muscle function, and complete knockout results in perinatal lethality ^22^. To investigate the role of *Ryr1 in vivo*, Lécuyer *et al.*, generated fetal liver chimeras in which hematopoietic cells lacked ryanodine receptor 1 (unpublished, manuscript enclosed). In this setting, Ca^2+^ signaling via ryanodine receptors played a relevant function in T cells in EAE *in vivo*. Fetal liver chimeras with hematopoietic cells deficient in *Ryr1* had decreased disease activity compared with wild type controls. A lower frequency of CD25⁺ and CD69⁺ T cells in the spinal cord in wild type fetal liver chimeras was associated with milder disease activity. Moreover, this phenotype was reproduced in conditional knockout *CD4^cre^Ryr1^fl/fl^* mice used in our study, demonstrating that calcium signaling via RYR1 in T cells is essential for tissue damage in the context of neuroinflammation.

In contrast, our findings reveal that although RYR1 plays a critical role in TCR-dependent Ca^2+^ signaling, it is dispensable for CD4⁺ T-cell function in intestinal inflammation. To investigate this, anti-CD3 antibody were either systemically administered *CD4^cre^Ryr1^fl/fl^* mice to induce a cytokine storm and acute intestinal inflammation or naïve CD4⁺ T cells from *CD4^cre^Ryr1^fl/fl^*mice were transferred to *Rag1*^-/-^ mice resulting in colitis. These findings may reflect a disease-specific role of ryanodine receptor–mediated Ca^2+^ signaling. Both intestinal models differ significantly from EAE, primarily in their mode of T-cell activation. In EAE, the response to a self-protein of the CNS is restricted by major obstacles. In contrast to the colitis models, few T cells in the periphery are specific for the MOG peptide used for induction of EAE, and these T cells are under the control of peripheral tolerance mechanisms. In addition, immune cells’ access to the CNS is limited by the blood-brain barrier. The extent of CNS inflammation is also not only determined by T-cell infiltration, but also by T-cell reactivation strength within the CNS^67^. In the anti-CD3 model, T-cell stimulation is polyclonal, leading to unspecific activation of T cells, including memory and effector T cells, regardless of their activation threshold and antigen specificity. This can override the physiological antigen presentation and co-stimulation process, where RYR1-mediated Ca^2+^ signaling might play a modulatory role. Similarly, in the T-cell transfer colitis, transferred T cells are exposed to a vast variety of microbial antigens of the intestine, which lead to pathogenic T-cell activation, in the absence of Treg-cell control. This diverse and high-antigen-load environment might be sufficiently potent to trigger T-cell activation in the absence of RYR1, thereby abrogating the requirement for RYR1-mediated Ca^2+^ release in T cells in this context. These factors suggest that EAE relies more on cell-intrinsic modulation of TCR signaling than the intestinal inflammation induced by anti-CD3 or adoptive T-cell transfer.

In conclusion, our findings suggest that RYR1 has a redundant function in naïve CD4⁺ T-cell activation, proliferation, and differentiation *in vitro*, as well as in both acute and chronic models of T-cell-mediated intestinal inflammation *in vivo*. These results contrast with the previously reported roles of RYR1 in neural inflammation, highlighting a potential disease-specific function of RYR1 in modulating immune responses. Therefore, further studies are warranted to uncover the underlying mechanisms explaining the different role of RYR1 signaling in CD4^+^ T cells in different diseases.

## Materials and Methods

### Mice

*Rag1^-/-^* mice were purchased from Jackson Laboratories. The following reporter mice were used: IFN-γ^Katushka^ ^37^, IL-17A^eGFP^ ^29^, and Foxp3^RFP^ ^38^. The mice were kept under specific pathogen free (SPF) conditions at the University Medical Center Hamburg-Eppendorf. All animal procedures were approved by the review board of the City of Hamburg (Behörde für Justiz und Verbraucherschutz der Freien und Hansestadt Hamburg). Age- and sex-matched littermates between 7 and 30 weeks were used.

### Generation of *Ryr1* conditional knockout strain

To generate a conditional *Ryr1* allele, we used a conditional ready (*Ryr1*^tm1a(EUCOMM)Hmgu^) construct from EUCOMM, in which exon 8 of the mouse *Ryr1* gene is flanked by loxP sites, ^39^. The L1L2_Bact_P cassette was inserted into the critical exon(s) (Build GRCm39). The cassette was composed of an FRT site followed by lacZ sequence and a loxP site. This first loxP site is followed by a neomycin resistance gene under the control of the human beta-actin promoter, SV40 polyA, a second FRT site and a second loxP site. A third loxP site is inserted downstream of exon 8 (Figure 1A). This entire construction functions as a selection cassette. To excise the selection cassette and generate a floxed *Ryr1* allele (*Ryr1*^tm1c(EUCOMM)Hmgu^), *Ryr1*^tm1a(EUCOMM)Hmgu^ mice were crossed with a Flp deleter line ^40^. The resulting conditional *Ryr1* knockout line *Ryr1*^tm1c(EUCOMM)Hmgu^ was then crossed with the CD4^cre^ (B6·Cg-Tg(Cd4-cre)1Cwi) line ^41^, referred to in the text as *CD4^cre^Ryr1^fl/fl^*.

### Isolation of spleen, lymph nodes, and intestinal leukocytes

Cells from the spleen and lymph nodes were collected by pressing the organs through 100 µm strainers, followed by filtration through 40 µm strainers. The colon was gently separated from the surrounding mesentery tissue, opened longitudinally, and washed in cold phosphate-buffered saline (PBS). It was then cut into small pieces and incubated two times in DTT buffer (containing 10% 10X Hank’s balanced salt solution, 10% 10X HEPES-bicarbonate buffer, 5% fetal bovine serum (FBS), and 5mM ethylenediaminetetraacetic acid (EDTA) in water) at 37°C for 30 minutes, with shaking. Next, the colon pieces were strained through a mesh into gentleMACS™ Dissociator C tubes (Miltenyi Biotec, Bergisch-Gladbach, Germany), in which intraepithelial lymphocytes (IELs) were collected by centrifugation, the supernatant was discarded, and the remaining colon pieces were incubated at 37°C for 30 min in collagenase solution (10% FBS, 1% 100x HGPG (HEPES, L-glutamine, penicillin/streptomycin, and gentamycin), 1 mM CaCl_2_, 1 mM MgCl_2_, 100 U/ml collagenase, 10 U/ml DNAse in RPMI 1640 medium). The C tubes were then attached upside down onto the sleeve of the gentleMACS Dissociator to obtain lamina propria lymphocytes (LPLs). The digested colon pieces were strained through a mesh strainer and added to the IEL fraction. LPLs and IELs were enriched by using a 67-40% (two-phase) Percoll gradient or 40% (one-phase) Percoll centrifugation through collecting the interphase or cell pellet, respectively.

### T-cell isolation

Total CD4^+^ cells were enriched using magnetic-activated cell sorting (MACS-Miltenyi Biotec) via positive selection, while naïve CD4^+^ T cells were enriched by negative selection using the EasySep^TM^ Mouse Naïve CD4^+^ T-cell isolation kit (STEMCELL Technologies, Vancouver, Canada) according to the manufacturers’ instructions. Cell purity exceeded 95% for total CD4^+^ cells and 86% for naïve CD4^+^ T cells measured by flow cytometry.

### *In vitro* CD4^+^ T-cell differentiation

CD4^+^ CD62L⁺CD44^low^ naïve T cells were sorted using FACS from enriched total CD4^+^ T cells (as described above). For each differentiation condition, the cells were cultured in a 96-well plate at 1×10^5^ or 2×10^5^ cells per well in 200 µL of full Click’s medium (10% FBS, 50 U/mL penicillin, 50 µg/mL streptomycin, 2 mM L-glutamine, and 0.05 mM β-mercaptoethanol) supplemented with the following cytokines and antibodies for 4 days. For differentiation of Th1 cells, naïve CD4^+^ T cells were cultured in the presence of 100 U/mL mIL-2, 10 ng/mL mIL-12 and 10 µg/mL anti-IL-4 mAb (clone: 11B11), and 2 µg/mL anti-CD28 mAb (clone: 37.51) in plates coated with 10 µg/mL anti-CD3 mAb (clone: 145-2C11). For the differentiation of Th17 cells, naïve CD4^+^ T cells were cultured in the presence of 10 ng/mL mIL-6 and 0.25 ng/mL hTGF-β1, 10 µg/mL anti-IL-4 mAb, 10 µg/mL anti-IFN-γ (clone: XMG1.2) mAb and 2 µg/mL anti-CD28 mAb in plates coated with 10 µg/mL anti-CD3 mAb. For differentiation of Treg cells, naïve CD4^+^ T cells were cultured in the presence of 50 U/mL mIL-2 and 2 ng/mL hTGF-β1 and 2 µg/mL anti-CD28 mAb in plates coated with 2 µg/mL anti-CD3 mAb. We obtained murine IL-2, IL-6, and IL-12 from Miltenyi Biotec and human TGF-β1 from BioLegend (San Diego, CA, USA). The anti-mouse CD3 mAb, anti-mouse CD28 mAb, anti-mouse IFN-γ mAb, and anti-mouse IL-4 mAb were produced in-house.

### *In vitro* CD4^+^ T-cell activation and proliferation

Naïve CD4^+^ T cells isolated using the EasySep^TM^ Mouse Naïve CD4^+^ T-cell isolation kit were stimulated with 2 µg/ml plate bound anti-CD3 and 1 µg/ml soluble anti-CD28 for 4 hours. T-cell activation was measured by flow cytometric analysis of NUR77 and IRF4. For measurement of T-cell proliferation, EasySep^TM^ -purified naïve CD4^+^ T cells were loaded with 2 mM CellTrace^TM^ violet stain (Invitrogen Thermo Fisher Scientific, Carlsbad, California, USA) and stimulated with 2 µg/ml plate-bound anti-CD3 and 1 µg/ml soluble anti-CD28 for 48 hours. Proliferation was determined by dye dilution via flow cytometry.

### Immunofluorescence staining

Immunofluorescence of RYRs in primary T cells were done according to ^42^. In brief, freshly isolated WT and *Ryr1^-/-^* naïve CD4^+^ T cells were seeded on slides coated with poly-l-lysine (0.1 mg/ml). The cells were fixed with 4% (w/v) para-formaldehyde for 15 min and permeabilized with 0.05% (v/v) saponin for 15 min. To block nonspecific binding sites, cells were incubated with 10% (v/v) FBS over night at 4°C. Primary mouse anti-RYR (1:100, 34C, GTX22868, GeneTex) was diluted in 3% (v/v) FBS and incubated over night at 4°C. The secondary antibody (anti-mouse Alexa Fluor 488, 1:400, A21202, Thermo Fisher Scientific) was diluted in 3% (v/v) fetal bovine serum and incubated for 1 hour at room temperature. For nuclear staining, the cells were incubated with 4′,6-diamidino-2-phenylindole (DAPI; 62247, Thermo Fisher Scientific, 1:1000) for 10 min at RT. Coverslips were mounted with Abberior Mount Solid (Abberior) overnight at 4°C. Images were acquired using a super-resolution spinning disc microscope (Visitron) equipped with a CSU-W1 SoRa (super-resolution via optical re-assignment) Optic (2.8×, Yokogawa), a 100× magnification objective (Zeiss), and a scientific complementary metal-oxide-semiconductor camera (Orca-Flash 4.0, C13440-20CU, Hamamatsu) in SoRa mode (280x). The following lasers and filters were used for the respective dyes and fluorophores: DAPI, excitation 405-nm laser and emission 460/50-nm filter; Alexa Fluor 488, excitation 488-nm laser and emission 525/50 nm).

### Flow cytometry

For extracellular staining, cells were resuspended in 100 µL PBS containing Live/dead staining dye (Pacific Orange™ succinimidyl ester, triethylammonium salt, Invitrogen Thermofisher) for 10 min at 4°C. After washing, cells were stained with fluorochrome-conjugated antibodies in PBS containing 1% FBS for 10 min at 4°C.

For intracellular staining, cells were restimulated for 4 hours with PMA (phorbol 12-myristate 13-acetate, 50 ng/ml, Sigma-Aldrich, St. Louis, MO, USA), ionomycin (1 mM, Sigma-Aldrich), and monensin (2 µM, BioLegend) for 4 hours at 37°C. After restimulation, extracellular staining was performed as described above. Then cells were fixed in 10% Formaldehyde for 10 min in the dark and at room temperature (RT), permeabilized with 0.1% NP-40 (5 minutes at RT and dark), and stained intracellularly for 1 hour.

The BD LSRFortessa and the FACSAria Fusion/Illu (BD Biosciences, San Jose, CA) were used for cell analysis and cell sorting, respectively.

Extracellular antibodies: anti-CD45 mAb (clone: I3/2.3, AF700), anti-CD3 mAb (clone: 17A2, BV421, BV650, BV785), anti-CD4 mAb (clone: GK1.5, BV785, clone: RM4-5, PE-Cy7), anti-CD8α mAb (clone: 53-6.7, BV650, PE-Cy7), anti-CD62L mAb (clone: MEL-14, APC, BV510), CD44 m Ab (clone: IM7, APC-Cy7).

Intracellular antibodies: anti-IL-17A mAb (clone: TC11-18H10.1, AF-488), anti-IL-10 mAb (clone: JES5-16E3, PE, PE-Cy7), anti-TNF-α mAb (clone: MP6-XT22, APC, BV421), anti-IFN-γ mAb (clone: XMG1.2, BV711, BV785), anti-NUR77 mAb (clone:12.14, PE), anti-IRF4 mAb (clon: 3E4, PE-Cy7). All the antibodies used were bought from Biolegend.

### Gating strategy

For every flow cytometry analysis, cells were first gated on FSC-A vs. SSC-A to select lymphocytes with moderate size (FSC-A) and low granularity (SSC-A). Singlets were then gated using FSC-A vs. FSC-H. For *ex vivo* analyzes, live CD45⁺ leukocytes were identified by excluding dead cells (PacOrange⁺) and gating on CD45⁺ cells, followed by selection of CD3⁺CD4⁺ T cells. For *in vitro* assays, viable cells were similarly identified by exclusion of PacOrange⁺ events, and CD4⁺ T cells were gated directly. A representative gating strategy for *ex vivo* flow cytometric analysis of colonic CD4⁺ T-cell subsets is shown in Fig. S1.

### Transfer colitis and colonoscopy

For the naïve T-cell transfer colitis, CD45^+^RB^high^CD4^+^ T cells were sorted using FACS, and a total of (200×10^3^) cells were injected intraperitoneally (i.p.) into *Rag1^-/-^* mice.

For the cytokine-polarized T-cell transfer model, total CD4^+^ T cells from *CD4^cre^Ryr1^+/+^ IFN-γ^katushka^ IL-17^eGFP^Foxp3^RFP^* (*CD4^cre^Ryr1^+/+^*) and *CD4^cre^Ryr1^fl/fl^ IFN-γ^katushka^ IL-17^eGFP^Foxp3^RFP^*(*CD4^cre^Ryr1^fl/fl^*) mice were polarized *in vitro* under Th17 conditions (10 ng/mL mIL-6, 0.25 ng/mL hTGF-β1, 10 ng/mL mIL-1β, 20 ng/mL mIL-23, 10 µg/mL anti-IL4 mAb, 10 µg/mL anti-IFN-γ mAb, and 2 µg/mL anti-CD28 mAb in plates coated with 10 µg/mL anti-CD3 mAb) and Th1 conditions (100 U/mL mIL-2, 10 ng/mL mIL-12 and 10 µg/mL anti-IL4 mAb, and 2 µg/mL anti-CD28 mAb in plates coated with 10 µg/mL anti-CD3 mAb). After 4 days, Th17 (IL-17A^eGFP+^) and Th1 (IFN-γ^Katushka+^) cells were sorted by FACS. Either Th17 cells alone (5×10^4^) or 1:1 mixture of Th17 with Th1 cells (each subset 2.5×10^4^) were injected i.p. into *Rag1^-/-^* mice. Mice were assessed weekly for colitis development and severity by endoscopy (custom-made colonoscopy system, Karl Storz, Tuttlingen, Germany) as previously described ^43,44^. Mice were anesthetized with isoflurane and scored based on stool consistency, granularity of mucosal surface, vascularity changes, and colon wall translucency. Each parameter was graded on a scale from 0 to 3, with an overall score ranging from 0 (healthy) to 12 (severe colitis).

### Anti-CD3 mAb-induced intestinal inflammation

Mice were injected i.p. with 15 µg anti-CD3 mAb (clone: 145-2C11) at the beginning of the experiment, followed by two injections at 48-hour intervals and sacrificed 4 hours after the third injection. Mice weight was measured every day.

### Quantitative RT-PCR

Total CD4^+^ T cells were isolated as described before from spleens and small intestine of *IFN-γ^katushka^ IL-17^eGFP^Foxp3^RFP^* triple reporter mice under steady-state condition or anti-CD3 mAb-induced inflammatory condition. For the steady state analysis, cells were sorted by FACS for naïve (CD62L⁺CD44^low^), effector (CD62L⁻CD44^high^), and regulatory (Foxp3^+^) CD4^+^ T cells. In the inflammation model, mice were treated with anti-CD3 mAb as described above, and 4 hours after the last injections, total CD4^+^ T cells were isolated from the small intestine. Th1 (IFN-γ^Katushka+^), Th17 (IL-17A^eGFP+^), and Th1/Th17 cells (IL-17A^eGFP+^ IFN-γ^Katushka+^), as well as Foxp3^RFP^⁺ Treg cells were sorted using FACS.

RNA was isolated using Trizol (Life Technologies, Carlsbad, CA, USA) and reverse transcribed using random primers. cDNA concentrations for *Ryr1* and *Hprt* were analyzed by TaqMan real time PCR using *Ryr1* and *Hprt* primers (Thermo Fisher Scientific, CA, USA). *Ryr1* values were normalized according to *Hprt* values.

### Measurement of localized Ca^2+^ signals

3-5×10^6^ EasySep^TM^ isolated naïve CD4^+^ T cells were loaded with Ca^2+^ indicators, 10 µM Fluo-4 AM and 20 µM FuraRed AM in RPMI1640 with 10% FCS for 20 minutes in the dark at RT. After 20 min, 2 ml of the same medium was added, and cells were incubated for an additional 30 min. After washing cells with Ca^2+^ measurement buffer (140 mM NaCl, 5 mM KCl, 1 mM MgSO_4_, 1 mM CaCl_2_, 20 mM HEPES (pH 7.4), 1 mM NaH_2_PO_4_, 5 mM glucose), cells were resuspended in the same buffer. For imaging, 10 µL of cells were added on coverslips precoated with bovine serum albumin (5 mg/mL, Sigma-Aldrich) and poly-L-lysine (0.1 mg/mL, Sigma-Aldrich) which promotes cell adhesion and minimizes movement during measurement. G magnetic beads (10 µm in diameter, Merck Millipore, Burlington, Massachusetts, USA) were coated with 0.5 mg/ml anti-CD3 mAb (clone 145-2C11, BD Pharmingen™ BD Biosciences, San Jose, CA, USA) and 0.5 mg/ml anti-CD28 mAb (clone: 37.51, BD Pharmingen™ BD Biosciences) for 30-60 minutes at RT with rotation, then washed and suspended in Ca^2+^ measurement buffer. Imaging was performed using a 100x oil immersion lens on a brightfield light microscope with a xenon arc lamp as a light source and a dual-view module for both Fluo-4 and FuraRed signals. After measuring the basal activity for 1 minute, anti-CD3 and anti-CD28-coated beads were directly added to the imaging field without disturbing the slide. Ca^2+^ responses were recorded for an additional 2 min. Ca^2+^ microdomains were measured and analyzed as described in ^45^.

### Measurement of global Ca^2+^ signals

An aliquot of 10^7^ freshly EasySep^TM^-isolated murine naïve CD4^+^ T cells were loaded with the membrane-permeable AM ester of Fura-2 (4 µM) for 15 min at 37°C in 1 mL of full Click’s medium. After 15 min, 4 mL of fresh medium was added. The cells were rinsed twice and resuspended in Ca^2+^ measurement buffer. Cells were added on prepared coverslips, coated with bovine serum albumin (5 mg/mL, Sigma-Aldrich) and poly-L-lysine (0.1 mg/mL, Sigma-Aldrich).

Cells were stimulated by 2 µg/mL anti-CD3 mAb (clone 145-2C11, BD Pharmingen™ BD Biosciences) after 20 s of recording. Imaging was performed on a Leica IRBEmicroscope (Wetzlar, Germany) with 40x-fold magnification. A Sutter DG-4 (Novato, California, USA) was used as a light source, and frames were acquired with an electron-multiplying charge-coupled device camera (Hamamatsu, Hamamatsu City, Shizuoka Prefecture, Japan). One frame every two seconds was acquired using a Fura-2 filter set (excitation, HC 340/26, HC387/11; beam splitter, 400DCLP; emission, 510/84; all in nanometers; AHF Analysentechnik (Tübingen-Pfrondorf, Germany)). Intracellular Ca^2+^ concentration [Ca^2+^] _i_ was determined in Fura-2 loaded murine CD4^+^ T cells. Therefore, R_min_ [using the lowest ratio (R) and fluorescence (F) after EGTA chelation] and R_max_ [using the highest R and F after Ionomycin incubation] of Fura-2 in single-cell measurements were assessed.

### Statistics

Statistical analyses were performed using GraphPad Prism (GraphPad Software, La Jolla, CA), and flow cytometry data were analyzed with FlowJo software (BD Biosciences). Statistical tests used are specified in the figure legends. P-value of p < 0.05, p < 0.01 and p < 0.001 were considered statistically significant and are denoted as *, ** and ***, respectively.

### Experimental schematics

Schematic illustrations were created using BioRender.com.

## Data Availability Statement

The data supporting the findings of this study are available from the corresponding author upon reasonable request.

## Ethics Statement

All animal experiments were approved by the review board of the City of Hamburg (Behörde für Justiz und Verbraucherschutz der Freien und Hansestadt Hamburg; Registration Nos. N 67/20 and N 021/25). No human studies were performed.

## Authors contribution

Samuel Huber, Hans-Willi Mittrücker, Mikolaj Nawrocki, and Tanja Bedke conceived and supervised the study. Sogol Dostiar Tabrizi and Mikolaj Nawrocki performed experiments, analyzed the data, and wrote the manuscript. Björn-Phillip Diercks and Franziska Möckl provided expertise in Ca²⁺ imaging and data analysis. Lola Hernandez carried out the immunofluorescence staining. Friederike Stumme, Melina Birus, and Marius Böttcher assisted with experimental processing. Andreas Guse, Nicola Gagliani, Dmitri Lodygin, and Alexander Flügel provided critical input and feedback on the manuscript. All authors reviewed and approved the final version.

## Acknowledgments

We thank Dr. Morsal Sabihi for manuscript editing, Cathleen Haueis for technical assistance, and the UKE FACS Core Facility for their support with cell sorting.

