## Supplementary Figure 1 for "Ryanodine receptor 1 is dispensable for CD4^+^ T-cell differentiation and effector function in intestinal inflammation models"

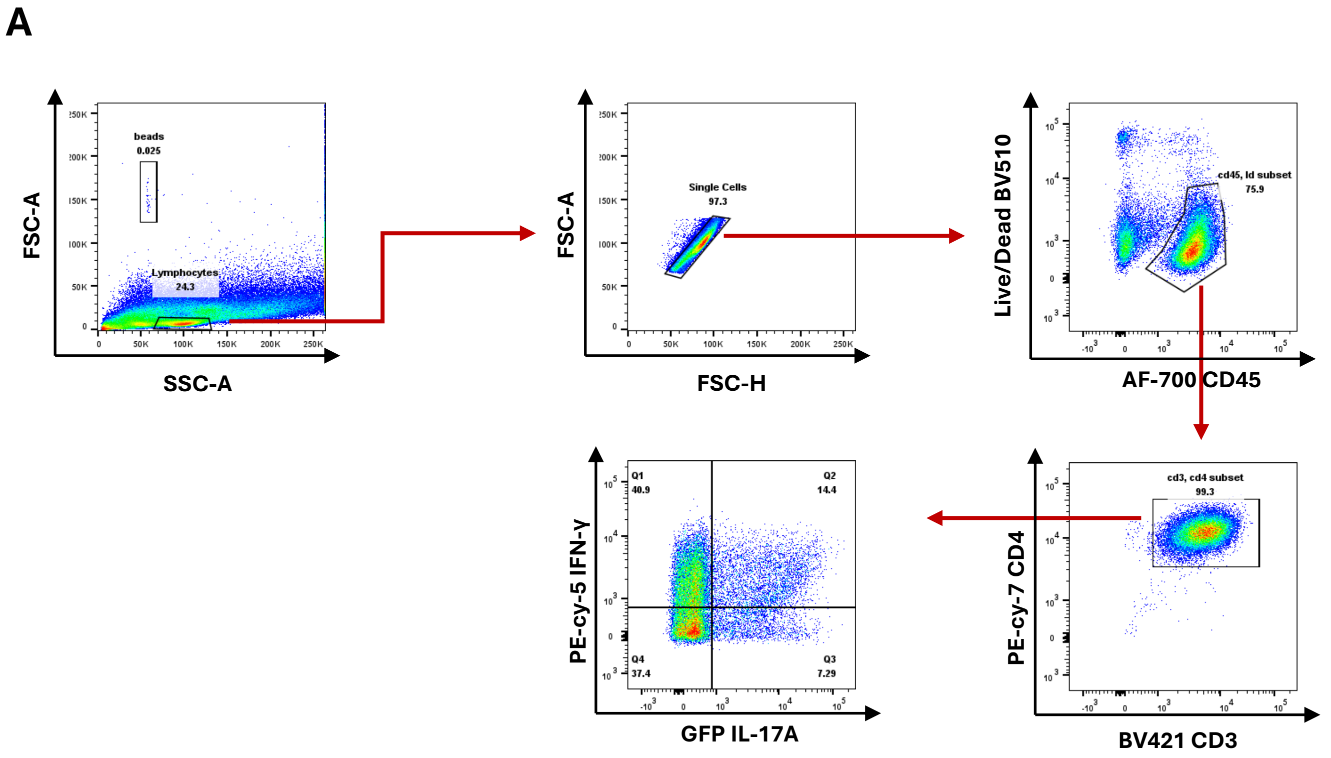


**Fig. S1. Gating strategy for identification of CD4⁺ T cell subsets from the colon.**
**A)** Representative flow cytometry plots showing the sequential gating strategy used for *ex vivo* analysis in the Th17 adoptive transfer colitis model (corresponding to Fig. 7). Debris was excluded based on FSC-A and SSC-A characteristics, and single cells were identified by gating on FSC-A and FSC-H. Live $CD45^+$ leukocytes were gated using a viability dye (Live/Dead BV510 vs. AF700 CD45). From this population, $CD3^+CD4^+$ T cells were identified. Within the $CD4^+$ T cell population, IL-$17A^+$ ($GFP^+$) and IFN-$\gamma^+$ (PE-$Cy5^+$) subsets were quantified.
All frequencies and absolute numbers were calculated relative to the total $CD4^+$ T cell population. Representative staining is shown.
